# Auditory cortical population codes for pitch are cue dependent

**DOI:** 10.64898/2026.11.25.769094

**Authors:** Veronica M Tarka, Quentin Gaucher, Andrew J King, Ben DB Willmore, Kerry MM Walker

## Abstract

Pitch provides a powerful example of perceptual invariance, enabling us to recognize the same melody across different instruments. The fundamental frequency (F0) at which a pitch is heard can be derived from either the harmonic spacing of frequency components (“spectral” cues), the waveform periodicity (“temporal” cues), or both. It is unclear whether population codes for F0 in auditory cortex are invariant across these acoustic cue classes. We addressed this question by training a linear decoder to predict F0 from the spiking responses of 1085 neurons in ferret primary auditory cortex to 10 sound types varying in their spectral and temporal composition. F0 could be decoded accurately when trained and tested on any individual sound type. However, decoding performance generalised well across sounds only if they contained the same class of pitch cue, and poorly between complex sounds and pure tones. Onset and offset responses carried different information about the F0 of complex sounds. Finally, approximately 20 neurons were sufficient to decode F0, but F0 information was widely distributed across the neural population. These findings show that F0 is represented in primary auditory cortex in cue-dependent, temporally multiplexed subspaces, with sparse pitch representations embedded in a distributed, robust population code.

## Introduction

The tonal (“low” to “high”) perceptual quality of sound, known as pitch, gives rise to our experience of melody in music and prosody in speech, and helps us to focus on a single voice in a crowded room^1–4^. Pitch perception is often degraded by ageing and hearing loss^5,6^, and is highly impoverished for cochlear implant users^7^. Therefore, understanding the cortical mechanisms for pitch processing is a clinical priority, in addition to a central question in sensory neuroscience.

Almost all natural periodic sounds are composed of many frequencies, yet we experience their pitch as a single fundamental frequency (F0). A core challenge for the auditory system is therefore to represent a common F0 across sounds whose frequency spectra and waveforms differ substantially, for example when different musical instruments (that differ in *timbre*) play the same note. The dual processing theory of pitch suggests that F0 is extracted from two distinct classes of acoustical cues^8^. The frequency components of a pitch-evoking sound are integer multiples (*harmonics*) of F0. Lower harmonics are “resolved” in the auditory nerve because the spacing between these harmonics is sufficiently large relative to the frequency tuning width of auditory nerve fibres to allow F0 to be represented in a population place code^9^. Higher harmonics are “unresolved”, but auditory nerve responses phase lock to the sound’s envelope periodicity, providing a temporal spiking cue for F0^10^. Because both spectral and temporal cues can support pitch perception^11–13^, the higher auditory system could in principle combine these representations to derive a cue-invariant representation of pitch.

The auditory cortex is a likely candidate for the site of pitch processing, as it is required for pitch judgements in humans^14–16^ and animals^17^. Auditory cortex contains specialized neurons that respond selectively to F0 derived from either spectral or temporal cues, with a small number of neurons that can generalize F0 across cue classes^18–21^. At the same time, the F0 of a given sound type is readily represented by broader populations of auditory cortical neurons, but these representations are also sensitive to sound timbre^22–30^. As most studies of F0 population coding have used only one sound type, it is not clear to what extent cortical population codes are able to provide an invariant representation of pitch across timbres and pitch cue classes.

At least three population-level representations of F0 are possible (Figure 1). Auditory cortex may provide a timbre-invariant F0 representation, such that the pitch of any harmonic sound source can be represented by a single population code (Figure 1a). Alternatively, cortical populations may retain the provenance of F0 cues, producing pitch representations that generalize within spectral or temporal cue classes, but not across them (Figure 1b). Finally, F0 representations may be timbre-specific, where different timbres (e.g. different musical instruments) activate different neural populations to represent the same pitch (Figure 1c). F0 could therefore be represented within a single neural subspace, two cue-specific subspaces, or multiple timbre-specific subspaces^31^.

**Figure 1.**
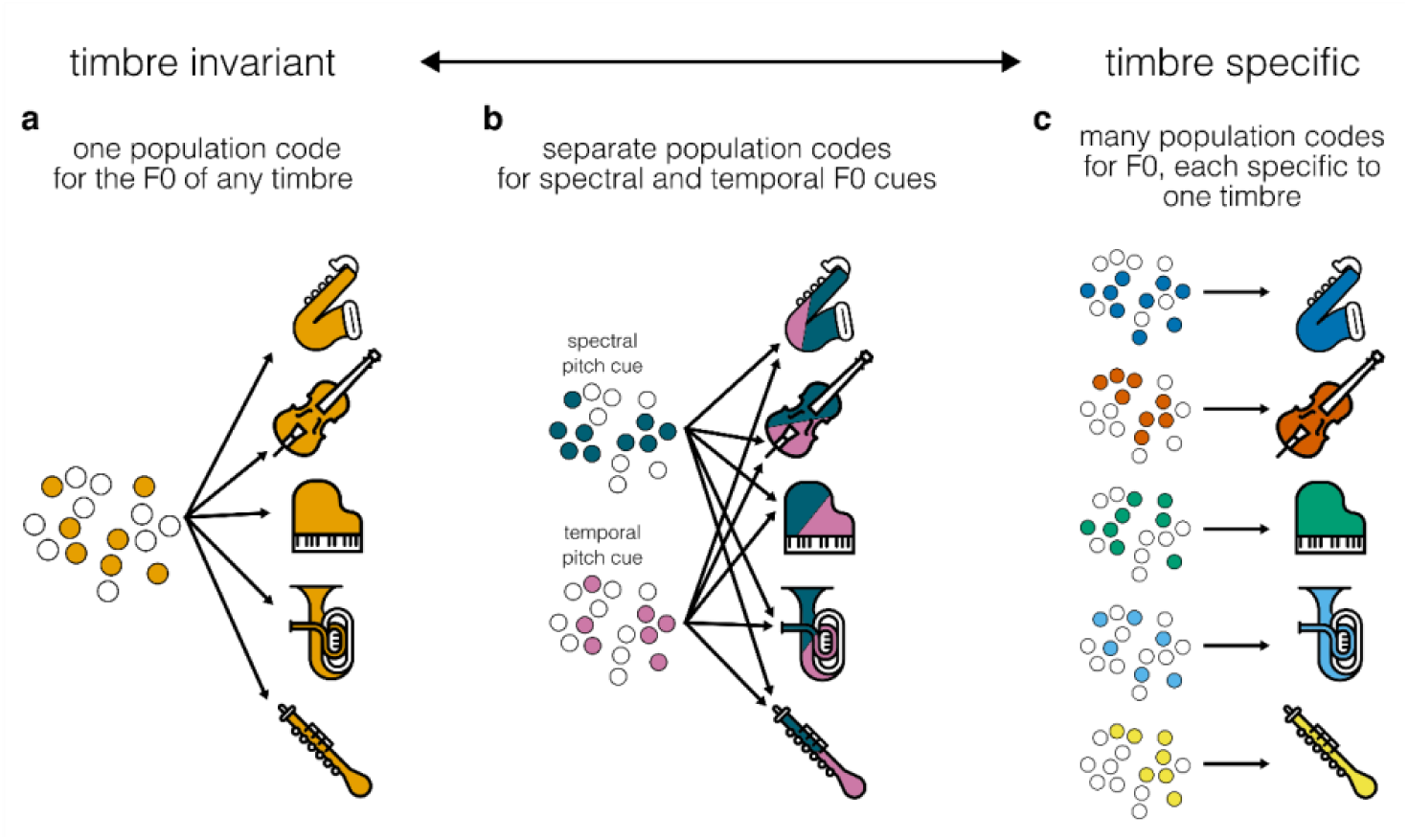
Schematic representations of three potential strategies for encoding F0 in auditory cortex. Populations of auditory cortical neurons are represented as circles, with filled circles indicating active neurons. These populations may provide: (**a**) a single representation of pitch that generalizes across all sound timbres; (**b**) two separate representations for spectral and temporal F0 cues; or (**c**) distinct F0 representations for every timbre.

We tested these competing hypotheses by presenting 10 stimulus timbres containing different combinations of spectral and temporal F0 cues while recording the spiking responses of 1085 neurons in the primary auditory cortex of anaesthetised ferrets. We trained linear decoders to predict F0 within each timbre and then tested how well these population codes generalised across timbres, response time windows, and neural subpopulations. We found that F0 can be decoded from population-level responses within every tested timbre, but generalization across timbres was strongest among sounds with the same F0 cue composition. Additionally, we found that population codes change throughout the neural response duration, with onset and offset responses providing different F0 information. The activity of a small number (∼20) of highly weighted neurons was sufficient to decode F0 nearly as well as the full population, even though the larger population could provide a robust F0 code in the absence of these neurons. These findings reveal cue-dependent and temporally multiplexed population codes for F0 in primary auditory cortex, rather than a single invariant pitch representation.

## Results

We examined how population codes in primary auditory cortex represent the F0 of stimuli across a range of timbres. We recorded the activity of 1266 well-isolated single units (“neurons”) from 4 ferrets (1077 in A1, 189 in AAF) to harmonic sounds that varied over 10 timbres (Methods; Figure 6) and 15 fundamental frequencies (in quarter octave steps from 354–4000 Hz). This included 633 (50%) neurons that were sensitive to both sound onset and offset, 234 (18%) sensitive to sound onset only, and 191 (15%) sensitive to sound offset only (paired t-tests, p<0.05). We trained a linear model to decode the F0 of the sound presented on each trial from the neural responses of these 1085 sound-responsive neurons (Figure 2a).

**Figure 2.**
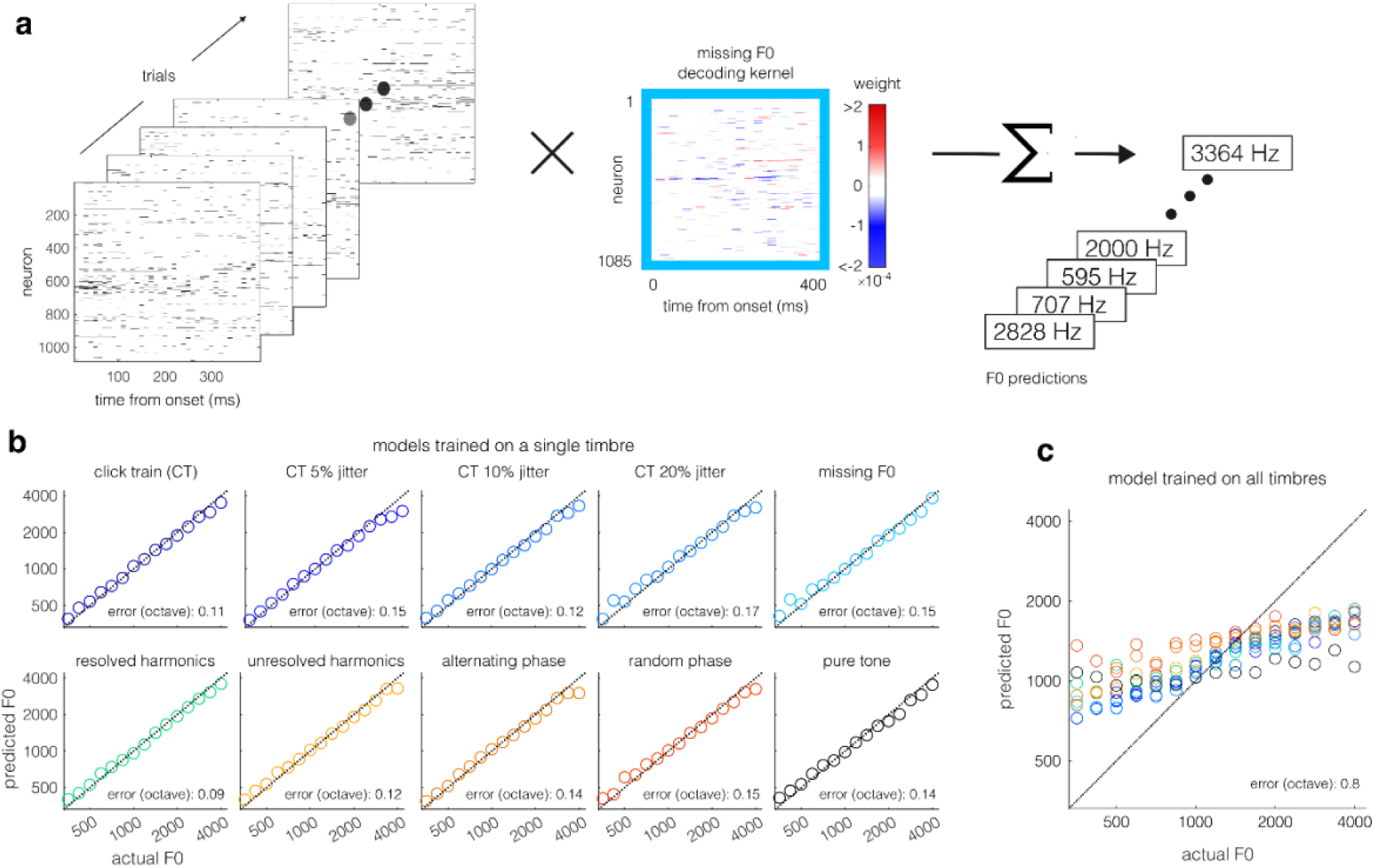
F0 can be linearly decoded from the cortical population. **a**) We decoded F0 on a trial-by-trial basis from the spiking responses of 1085 neurons. Spike counts on each trial were summed in 10 ms bins, taken from the onset of the sound until 200 ms following sound offset (400 ms total). A separate decoding model was trained for each of the 10 timbres presented. **b**) Each plot shows the performance of the decoding model on neural responses averaged across all trials with the same F0 for a single timbre. Colours correspond to pitch cue classes described in Figure 6. **c**) Performance of the best iteration of the model trained on neural responses to all timbres and tested on trial-averaged responses to each timbre as in **b**. Circle colours correspond to timbres shown in **b**. All errors reported are RMSE in octaves.

We began by training the decoder on trials within a timbre category. We found that F0 was decodable within all 10 timbres tested. F0 predictions on trial-averaged neural responses were on average 0.11 octaves from the true F0, corresponding to approximately 1 semitone on the Western musical scale (Figure 2b), with consistently high performance across 5 separate training folds (Figure S1a). Performance was best for the resolved harmonic timbre (Figure 2b, bottom left; 0.09 octave RMSE) and poorest for click trains with 20% jitter (Figure 2b, 2^nd^ to top right; 0.17 octave RMSE), reflecting the reduced perceived pitch salience with jittered click trains^32^. Manipulating the phase structure of unresolved harmonics disrupts their periodicity, leading to poorer pitch discrimination performance in ferrets^33^. This decreased pitch salience was also associated with relatively poor F0 decoding performance for high harmonics with randomized phase in the present analyses (Figure 2b; 0.12 to 0.15 octave RMSE). Our models decoded the F0 of single trials with an average RMS error of 0.55 octaves (Figure S1b). As expected, the accuracy is lower than for trial-averaged responses; this reflects trial-to-trial variability in neuronal responses. These initial analyses demonstrate that the population of primary auditory cortical neurons represents the F0 of harmonic sounds within a given sound timbre, allowing F0 classification even on individual trials.

To investigate whether a single population code could represent F0 invariantly across timbres, we next trained a single decoder to predict F0 using responses randomly drawn from all 10 timbres. The performance of this timbre-invariant model was substantially poorer than the timbre-specific models, with an RMSE of 0.8 octaves when tested on trial-averaged responses to each timbre (Figure 2c). Therefore, population F0 codes are not entirely timbre-invariant (Figure 1a).

### F0 population codes generalize within, but not across, pitch cue classes

To further examine the generalization of cortical F0 representations, we investigated whether these codes may be entirely timbre-specific (Figure 1c), or if they may instead encode F0 in terms of their acoustical pitch cue classes (Figure 1b). We used a decoding “transfer test”, which examined whether a F0 decoder trained on one timbre could reliably read out the F0 for a second, untrained timbre category (Figure 3a). Good generalization performance between any two timbres would indicate that they share similar population codes for F0.

**Figure 3.**
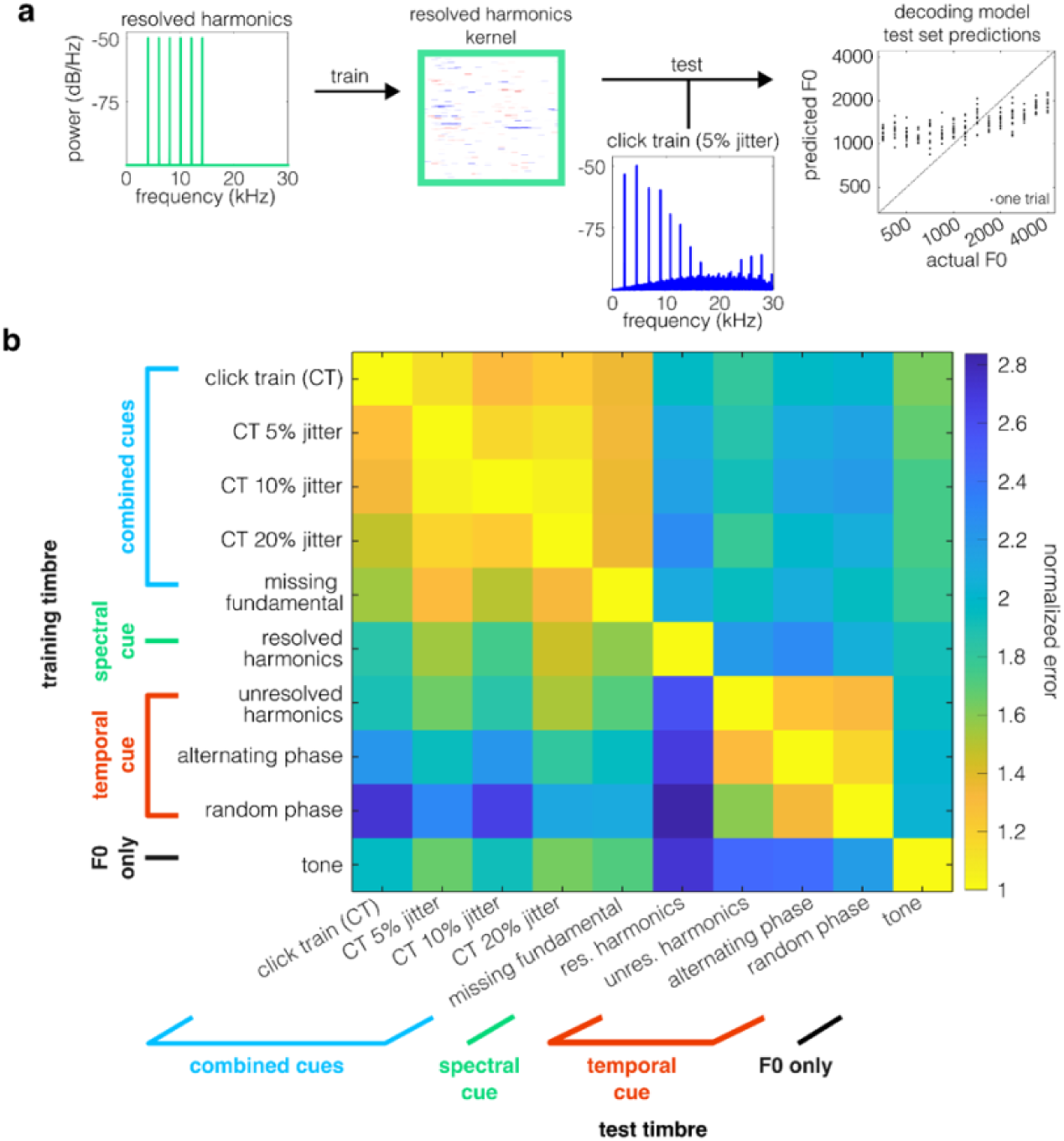
Different pitch cue classes are represented by distinct population codes. **a**) Schematic of an example cross-timbre F0 prediction. In this example, the F0 decoding model was trained on the resolved harmonics timbre (“training stimulus”), as described in Figure 2a. The fitted model was then used to decode the F0 of click trains with 5% jitter (“testing stimulus”). Accurate cross-timbre predictions would indicate that the neural population represented F0 similarly in the training and testing stimuli. **b**) Cross-timbre F0 predictions for all possible pairings of training (y-axis) and testing (x-axis) stimuli. Decoding accuracy (colour scale) was measured as the root mean squared error (RMSE) in octaves across trials, and normalized by the RMSE of the training timbre tested on itself (i.e. RMSE_cross-timbre_ / RMSE_within-timbre_). Lower values (yellow) indicate better cross-timbre decoding performance and therefore more similar F0 population codes between training and testing stimuli, while greater errors (blue) indicate poorer cross-timbre performance. Timbres that share F0 representations in the neural population clustered into four blocks of yellow values along the diagonal, corresponding to the four classes of pitch cues: resolved harmonics (spectral cues), unresolved harmonics (temporal cues), a combination of spectral and temporal cues, and pure tones.

We found that the matrix of decoding errors across all cross-timbre pairings was structured into distinct blocks, visible as yellow squares in the decoding matrix in Figure 3b. This suggests that the population F0 code generalizes better between some stimulus pairs than others. More specifically, cross-timbre generalization was generally high for stimulus pairs containing the same class of pitch cues. For example, decoders trained on a broad spectrum sound containing both resolved and unresolved harmonics could predict the F0 of other stimuli containing combined spectral and temporal cues with high accuracy, but not those containing only one of these pitch cues. Similarly, unresolved harmonic decoder weights could be used to predict the F0 of other sounds containing only unresolved harmonics, but performed poorly for resolved harmonic stimuli. Decoders trained on unresolved harmonics with randomized phase produced particularly poor F0 predictions for other timbres, consistent with the important role of envelope periodicity in the temporal F0 code. Pure tone training also generalized poorly to every other timbre tested, suggesting that there are separate population codes for representing pure tone frequency and the F0 of complex sounds. Together, these results indicate that F0 representations within populations of primary auditory cortical neurons can generalize F0 well within, but not across, temporal and resolved harmonic pitch cue classes. These population representations retain information about the acoustical cue type, rather than expressing a single cue-invariant pitch code.

### Cue-dependent F0 information is present in both onset and offset responses

The decoder weighted the response of each neuron in each 10-ms time bin (Figure 2a), allowing us to ask which time points in the neural response contained F0 information. We used principal component analysis (PCA) to investigate whether large-scale trends exist in the temporal structure of these weights, and whether these differed across pitch cue classes. We found that the strongest components captured broad onset and offset activity, rather than fine temporal spiking patterns (Figure 4a).

**Figure 4.**
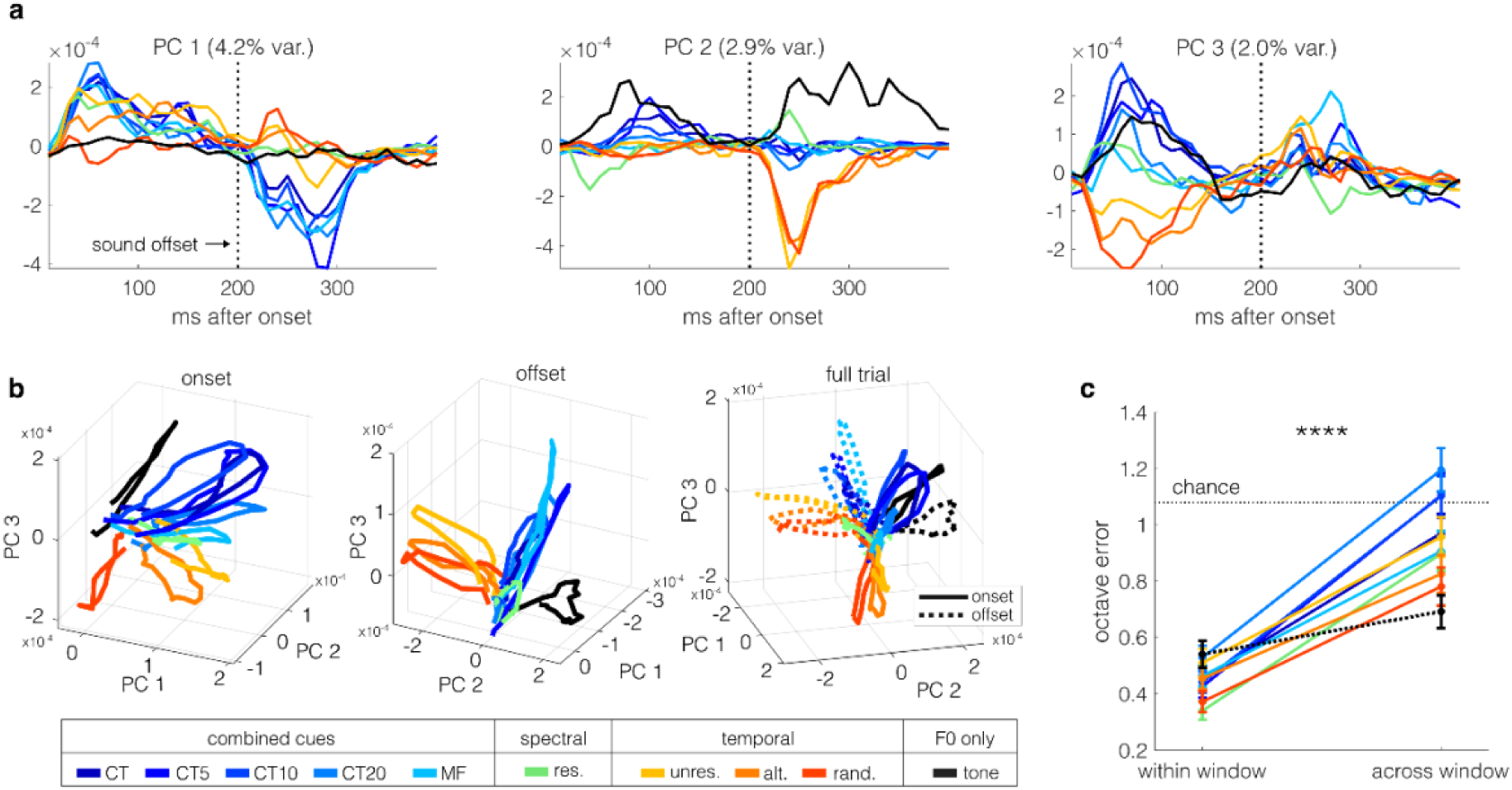
Onset and offset responses provide non-redundant, cue-dependent information about the F0 of complex sounds. **a**) Time course of the first three principal components (PCs; left to right) of the F0 decoding kernels. Colours indicate pitch cue classes (legend). **b**) Decoder kernel trajectories are plotted in the first three principal component space. Trajectories are plotted separated for trial onset (0-200 ms after sound onset, left panel), offset (0-200 ms after sound offset, middle panel), and the full trial (0-400 ms after sound onset, right panel). Colours as in **a**. These trajectories provide a low-dimensional visualization of the cue class grouping and the separation of onset and offset response weighting. **c**) Performance (mean octave error) for decoders trained and tested within the same response window or transferred across onset and offset windows. Across window decoding errors were significantly higher than within window errors (2-way ANOVA, ****p<0.0001) for every complex timbre tested (Tukey-Kramer HSD test, p<0.001, solid lines), but performance did not differ for pure tones (Tukey-Kramer HSD test, p>0.05, dashed line), indicating that onset and offset responses contain non-redundant information about the F0 of complex sounds.

The first three components explained 9.2% of the total kernel variance, with the scree plot showing a clear drop after 5 components (Figure S2). Although ∼9% is a limited fraction of the overall variance, clear patterns emerged, including a categorical separation of temporal patterns across timbre classes. The first principal component primarily captured offset responses to broad spectrum timbres containing combined F0 cues (Figure 4a, left), while the second component largely captured onset responses to tones and offset responses to sounds containing only unresolved harmonics (Figure 4a, middle). The third principal component characterised a gradient in onset responses that differed between tones, stimuli containing only unresolved harmonics, and those containing resolved harmonics (Figure 4a, right). In general, timbres with shared pitch cue classes had similar principal component projections during sound onsets and offsets (Figure 4b), indicating that the cue dependence of F0 codes was not confined to a single response window.

We further examined the temporal organization of population F0 codes by re-training the within-timbre F0 decoder using time windows that varied from 10 to 200 ms (Figure S3). We found remarkably stable decoding performance across these response window resolutions, supporting a coarse temporal organization of F0 codes into spike rates during an onset and offset window.

### Onset and offset responses provide non-redundant F0 information

Onset and offset projections for a given pitch cue class followed different, even orthogonal, trajectories in principal component space (Figure 4b, right)—particularly for complex sounds. This separation could indicate distinct, non-redundant F0 readouts across onset and offset responses. Alternatively, it could arise from a simple transformation such as complementary onset and rebound-offset tuning^34^. We tested these two explanations directly with a cross-window decoding transfer test.

If offset F0 tuning is simply the inverse of onset tuning due to rebound release from inhibition, the offset responses would contain little additional information for decoding F0. For each timbre, we trained the F0 decoder on the spike count responses in a single 200 ms time window starting either at sound onset or offset. The fitted decoder was then tested on held-out responses either from the same (“within window”) or other (“across window”) time window (Figure 4c). Performance was poorer for across window decoding (2-way ANOVA, F(1,1540) = 371.0, p = 3.08×10^-74^). The disadvantage for across window performance was observed for every complex timbre tested (Tukey-Kramer HSD test, p<0.001), but not for pure tones (Tukey-Kramer HSD test, p = 0.94). Thus, onset and offset responses carry non-redundant information about the F0 of complex sounds, whereas pure tone frequency decoding is consistent with simple rebound offset responses.

### F0 codes for different cue classes are built from partly overlapping neuronal ensembles

The above analyses show that different population codes represent the F0 of different pitch cue classes. We next aimed to identify the source of this code diversity. The F0 of spectral and temporal pitch cues may be carried by distinct subpopulations of neurons, or by overlapping groups of neurons whose F0 tuning changes for different timbres.

To distinguish between these possibilities, we quantified how much each neuron contributes to the population decoding performance. For each timbre, ∼10% of neurons held 50% of the total decoding weights in the population. We ranked neurons for each timbre by the absolute value of their fitted decoder weights, summed across time bins. Neurons in the top 10% of this ranking were considered highly weighted (n = 109 neurons per timbre condition), and we examined the degree to which these sets of neurons overlapped across timbres.

Stimuli that were more acoustically similar shared a larger proportion of highly informative neurons. For example, approximately 46% (50 of 109) of the highly informative neurons were shared across click trains that were unjittered and 5% jittered, but click trains and pure tones shared only 25% (27 of 109) of their highly informative neurons. This indicates that readouts are not built from a single group of F0-tuned neurons; instead, certain timbres share more highly weighted neurons than others.

The number of highly weighted neurons shared between timbre pairs is visualized as a matrix in Figure 5a, to facilitate comparison to the F0 generalization matrix in Figure 3b. Sounds with combined spectral and temporal pitch cues shared many highly weighted neurons with one another, and also have considerable overlap with stimuli containing only one of these cues (Figure 5a). In contrast, the pure-tone ensembles shared few high-weight neurons with other timbres. Therefore, cue-dependent decoding is not carried out by 4 unique pools of neurons. Rather, the same neurons can participate in multiple F0 codes for complex sounds.

**Figure 5.**
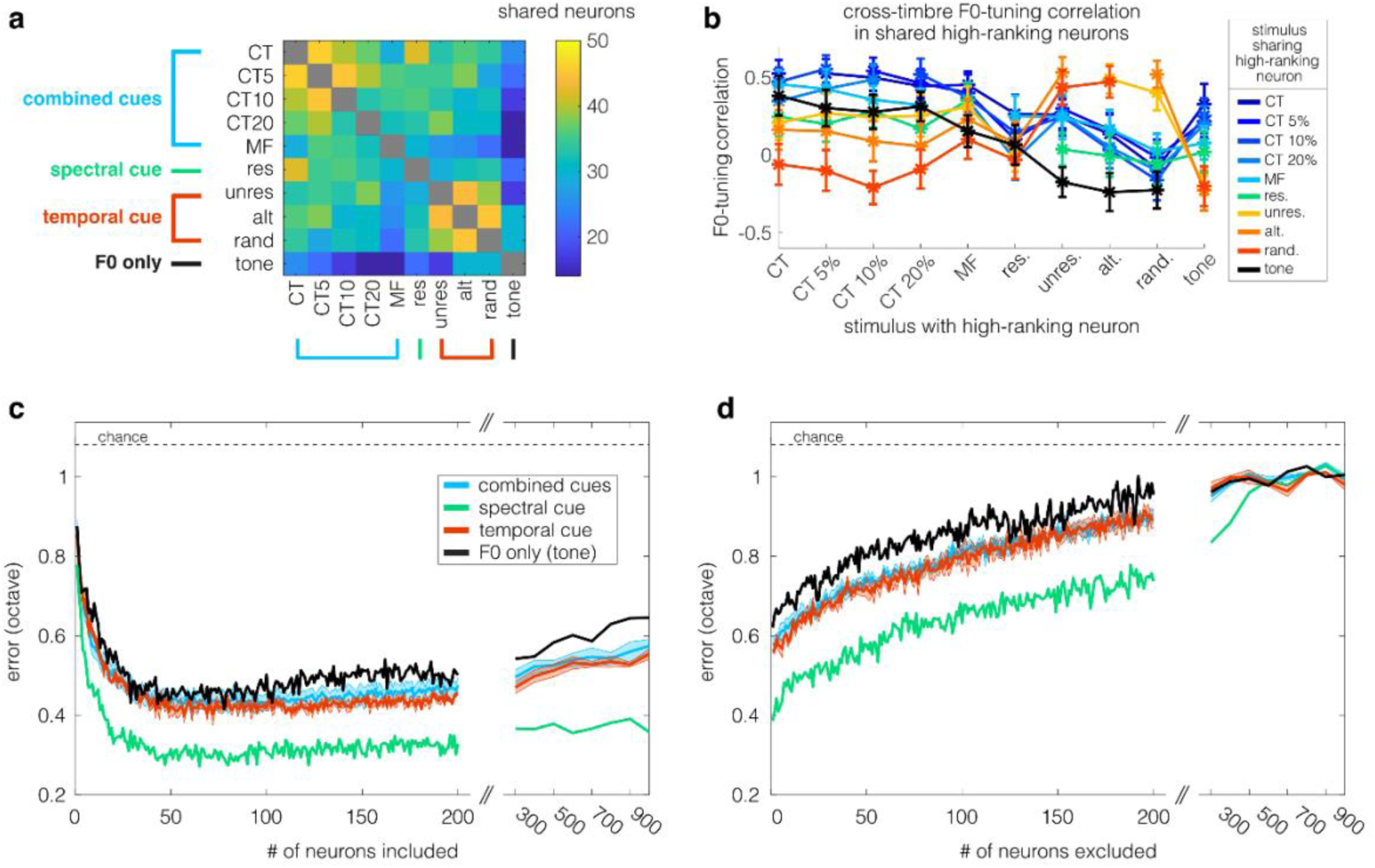
F0 representations are cue dependent and involve sparse, partially overlapping ensembles of neurons. **a**) Number of shared neurons with the top 10% (109) of F0 decoder weights for pairs of timbres. Overlap of neurons is greatest for stimuli that share spectral and temporal pitch cues. **b**) The mean correlation of F0 tuning curves across shared highly weighted neurons is shown for pairs of timbres. Tuning is more similar within pitch cue classes than across classes. Coloured lines indicate timbres. **c**) F0 decoding error as neurons are added to the model, in order of largest to smallest decoder weight (in the full model). Error steadily rises from the minimum achieved with ∼75 neurons due to the increasing number of model weights to be fit for decreasingly informative neurons. **d**) Decoding error as neurons are removed from the pool, from largest to smallest decoder weight. A small subset of neurons is sufficient for near-maximal decoding performance (**c**), but the remaining neurons support robust above-chance F0 decoding after these strongest contributors are removed (**d**). Error in both (**c**) and (**d**) was averaged across all timbres within a cue class (shown by line colour), and shaded error bars show standard error of the mean.

Consistent with this interpretation, we found that shared high-weight neurons had more similar F0 tuning across timbres in the same pitch cue class (Figure 5b). Neurons with large weights for pure tone frequency decoding showed similar tuning for the F0 of complex sounds only if there was energy present at F0 (i.e. click trains). Together, these results show that the differences in F0 codes for different pitch cue classes reflects *both* cue-dependent F0 tuning changes and contributions from different subsets of neurons.

### Sparse populations of neurons are sufficient, but not necessary, for F0 readout

As the above analysis showed that a small proportion of neurons had the highest weights in the decoder (Figure S4), we next asked how many of these high-weight neurons were sufficient to support F0 readout. Adding an increasing number of neurons to the pool in rank order showed that approximately 20 of the most highly weighted neurons were sufficient to decode F0 at 90% of the accuracy of the entire population (Figure 5c). The population code for F0 in primary auditory cortex is therefore sparse.

Finally, we asked whether F0 population decoding relies critically on this sparse subset of highly weighted neurons, or whether F0 could be reliably recovered from a large population of neurons with low decoder weights. We found that sufficiency did not imply necessity. When neurons were removed from the decoding pool in decreasing decoder weight order, decoding errors increased gradually and near linearly. Decoding remained better than chance even after the 200 most informative neurons had been removed (Figure 5d). This is particularly surprising given that half the decoding weights are held by the 100 highest ranked neurons (Figure S4).

These findings show that F0 information is concentrated enough for a few highly informative neurons to represent F0, but is still distributed enough to survive the loss of these neurons. This combination of sparse sufficiency and population-level robustness provides a powerful representation of F0 across the cortical population.

## Discussion

A defining property of pitch perception is constancy: sounds with very different spectra and temporal waveforms can evoke the same pitch. Here, we asked whether this perceptual invariance is reflected in a common population representation of fundamental frequency (F0) in primary auditory cortex. We linearly decoded F0 from the activity of 1085 auditory cortical neurons for a range of stimuli, and found that these codes were not invariant across all sound timbres. Instead, decoding generalized most successfully between sounds with similar combinations of spectral and temporal periodicity cues (Figure 3). This suggests that multiple population codes exist to represent F0 in a cue-dependent manner.

### Cue-dependent population representations of F0

Resolved harmonic structure and temporal periodicity provide complementary F0 information from which the same percept of pitch can be derived^9,13^. However, the existence of these pitch cue classes may not imply distinct neural codes. Leading theories of pitch encoding include spectral, temporal, and spectrotemporal representations at the cochlear level^35^. The relative importance of spectral and temporal cues to pitch perception differs across listening conditions^36^ and species (gerbils^37^, marmosets^38^, chinchillas^39^, and ferrets^33^).

Single neurons in ferret and marmoset auditory cortex can represent either spectral or temporal F0 cues^18,21^, with a small subset of neurons encoding both cues^19–21^. At the neural population level, resolved harmonics are represented as a place code in low-frequency regions of macaque A1, whereas neurons in higher frequency A1 can provide a spiking output that is temporally synchronized to the periodicity of low (<400Hz) F0 sounds^26,29^. Our results build on these findings to show that neural populations in ferret primary auditory cortex also provide distinct representations of F0 derived from spectral and temporal cues. Unlike the population codes described in macaques, the spike rate codes we identified do not require synchronisation to F0, and can therefore represent temporally-derived F0 for pitches above the phase-locking limit of auditory cortex.

Our results suggest neural mechanisms that explain why pitch representations in human auditory cortex are neither completely stimulus-specific nor fully generalized across timbres. Experiments using a range of pitch-evoking stimuli have often found distributed pitch-sensitive responses in the human auditory cortex rather than a single pitch-selective region^23,40^. Pitch-sensitive regions in anterior auditory cortex respond much more strongly to resolved than unresolved harmonics, paralleling the perceptual advantage of resolved harmonics in humans^41^. These results, together with the present findings, argue against treating pitch extraction as a process that fully integrates spectral and temporal cues at an early cortical stage.

Our findings suggest that, rather than fully integrating pitch cues, primary auditory cortex distils F0 information into a small number of low-dimensional, cue-dependent subspaces which depend on small, overlapping subsets of neurons and form building blocks for pitch representation. These could be the direct substrate for pitch perception themselves, but their low-dimensional nature means they could also be passed efficiently to downstream areas where they are integrated to form a universal representation of pitch. Furthermore, cortical representations may become more cue-invariant when the listener is actively listening to sounds. Indeed, in ferrets performing a pitch-discrimination task, auditory cortical activity was shown to carry information both about stimulus F0 and the animals’ reported pitch judgments^42^.

### F0 information in onset and offset responses

Our results show that neural responses to both sound onset and offset supported F0 decoding in a cue-dependent manner (Figure 4). For every complex sound tested, the population responses in these two time windows required different decoding weights to support accurate F0 categorization. In contrast, pure tone frequency decoding generalized between onset and offset responses. These results indicate that the offset population response to a complex sound is not simply a delayed or sign-reversed version of its onset F0 representation. Instead, the two windows provide at least partly non-redundant representations of the same perceptual feature.

Cortical offset responses are often tuned to frequencies that are non-overlapping with their onset best frequency^34,43–45^. Though offset responses have been previously thought to simply reflect a release from inhibition following an excitatory onset response, Scholl et al., 2010 demonstrated with intracellular recordings that onset and offset responses result from largely non-overlapping sets of synaptic inputs, which include both excitatory and inhibitory drives. Furthermore, separate on and off pathways are present in the auditory thalamus^46^, advancing theories that onset and offset responses arise through independent circuits in the auditory system^47^, as in the visual system.

Our findings support the claim that cortical offset responses are computationally richer than a passive rebound following sound termination. Cortical offset responses in individual neurons have been shown to represent multiple features of sound, including spatial location^48,49^, duration^50,51^, the order of sound sequences^52^, F0 and timbre^30^. Our findings extend this observation to population coding of F0. Cortical offset responses in mice were recently shown to be 3 times more accurate in decoding complex sounds than subcortical offset responses^53^. Onset and offset EEG responses in humans show distinct tuning properties, and are differently modulated by task demands^54^. Offset responses may also support the perceptual role that sound terminations play in sound segregation^55^, as they are associated with speech-in-noise perception in human listeners^56^.

The lack of linear transfer between onset and offset responses in our study demonstrates that auditory cortex can make F0 available through different population codes at different moments in time. Such temporal multiplexing has previously been described for F0 and timbre encoding in ferret auditory cortex^30^, and could allow downstream circuits to access different information about the pitch of a sound according to behavioural demands or sensory context^57^.

### Pure-tone frequency and the F0 of complex sounds

Pure tones evoke a salient percept of pitch, but they are the only sounds for which pitch is mapped along the cortical tonotopic axis. Thus, the F0 of pure tones can be extracted from this place code without further computation, whereas deriving F0 from complex sounds requires integration across frequency channels and/or throughout time. The frequency of pure tones was readily decoded from our cortical populations (Figure 2b), but this population code did not generalize to the F0 of any other timbre tested (Figure 3b). Furthermore, pure tone responses held more redundant F0 information across onset and offset windows than complex sounds (Figure 4c). These findings are consistent with distinct auditory cortical codes for pure tones and complex F0 in macaques^58,59^, humans^23,60,61^ and ferret single neurons^21,24^. This is compatible with the finding that only a small subset of auditory cortical neurons show similar F0 tuning for pure tones and complex harmonic sounds^20,21,23^, but the functional role of these specialized neurons in perception has yet to be tested.

### Sparse F0 codes are embedded within more distributed F0 representations

The F0 decoder heavily weighted a small proportion of neurons (Figure S4). In fact, approximately 20 of the most strongly weighted neurons were sufficient to achieve 90% of the F0 decoding accuracy obtained with the full population (Figure 5c). This agrees with previous work showing that decoding small ensembles of ferret auditory cortical neurons can match ferrets’ pitch discrimination for artificial vowels^24^. More generally, this fits within a wealth of studies showing that just tens of cortical neurons are sufficient for near-maximal discrimination of sensory stimuli (A1^62^, V1^63,64^) and motor responses (M1^65,66^). There was substantial overlap in the most F0-informative neurons across different timbres, especially within pitch cue classes. This suggests that the same neuron can participate in more than one F0 representation, but may show different temporal dynamics or tuning preferences for each. A downstream cortical area could therefore obtain accurate F0 estimates for all timbres explored here from a small cortical sample. An important caveat is that not all neurons in our population were simultaneously recorded, and so we shuffled trial order to remove noise correlations from those that were recorded simultaneously. Neural correlations can alter the information available in a neural population^67^, so the effects of noise correlations should be explored in larger-scale simultaneous recordings.

Even though a small number of neurons was enough to accurately represent pitch, we also found that pitch information was widely distributed across the cortical population (Figure 5d). This built-in redundancy across neurons would preserve F0 representation even if the most informative neurons were damaged or otherwise unavailable, and is consistent with previous evidence for distributed cortical representations of pitch^23,25,26^.

## Conclusions

Taken together, the present results show that spectral and temporal pitch cues recruit partially overlapping neuronal ensembles but distinct population codes in primary auditory cortex, with F0 information primarily in their onset and offset responses. Our results emphasize that F0 information for complex sounds is distinct from the representation of pure tone frequency. Pitch constancy observed in pitch-selective neurons^20,21^ may emerge from a downstream integration of these multiple cue-dependent representations of F0.

## Methods

### Animals

Data were collected from 4 adult ferrets (*Mustela putorius furo*; 2 female; age: µ = 29.3 weeks, σ = 18.4 weeks; Marshall BioResources, UK). All animal procedures in this study were approved by the local ethical review committee of the University of Oxford and performed under UK Home Office license.

### Surgical procedure

The dataset presented here were previously described as single neuron analyses in 2026^21^. Further details about the surgical procedures and stimuli are available in that publication. Briefly, terminal electrophysiological recordings were conducted under general anesthesia of ketamine (Vetalar; 5 mg/kg) and medetomidine (Domitor; 0.02 mg/kg). The use of anesthesia was employed to control for modulatory effects such as attention and arousal, and to yield stable recordings throughout cortical layers during long stimulus presentations. Animals were intubated and mechanically ventilated using medical-grade oxygen, and dexamethasone (0.5 mg/ml/hr) was provided to support respiration. Physiological parameters including blood oxygen levels, electrocardiogram activity, respiration rate, body temperature, and end-tidal CO₂ were continuously monitored. An ophthalmic ointment (Maxitrol; Alcon, UK) was applied to the eyes and atropine (Atrocare; 0.06 mg/kg) was administered periodically as required.

After securing the anaesthetized ferret in a custom-built stereotaxic apparatus, the shaved scalp was incised following bupivacaine injection (Marcain; <1 mg/kg), the temporal muscles were retracted, and a steel holding bar was affixed to the skull. A circular craniotomy (∼10 mm diameter) was made over the right auditory cortex, and the dura mater was removed. Stereotaxic coordinates (11 mm lateral to the midline and 8 mm anterior to Lambda) and the visible morphology of the ectosylvian gyrus were used to confirm the location of primary auditory cortex. The exposed cortical surface was covered with 1.25% agarose in 0.9% NaCl, and silicone oil was applied periodically between recordings.

The animal was transferred into an electrically and acoustically isolated anechoic chamber. A Neuropixels Phase 3^68^ microelectrode probe was advanced perpendicularly through the full depth of the auditory cortex. The location of each penetration within auditory cortex was determined using its visible position on the ectosylvian gyrus, local field potential latency, and frequency tuning. Of the 18 penetrations that produced sound-responsive activity, 10 were located in low-frequency A1 (primary auditory cortex), 3 in high-frequency A1, 2 in low-frequency AAF (anterior auditory field), and 3 in high-frequency AAF. We refer to all these regions collectively as primary auditory cortex. Neural data were recorded at a 30 kHz sampling rate using SpikeGLX software (https://github.com/billkarsh/SpikeGLX)^69^ and custom MATLAB scripts (MathWorks, Inc.).

### Sound stimuli

We presented 10 sound types (“*timbres”*), each over a range of 15 F0s (354 - 4000Hz in ¼ octave increments). Nine of the 10 timbres were complex sounds that included one of 3 classes of pitch cue: spectral cues, temporal cues, or combined spectral and temporal cues (Figure 6). The final timbre was pure tones.

**Figure 6:**
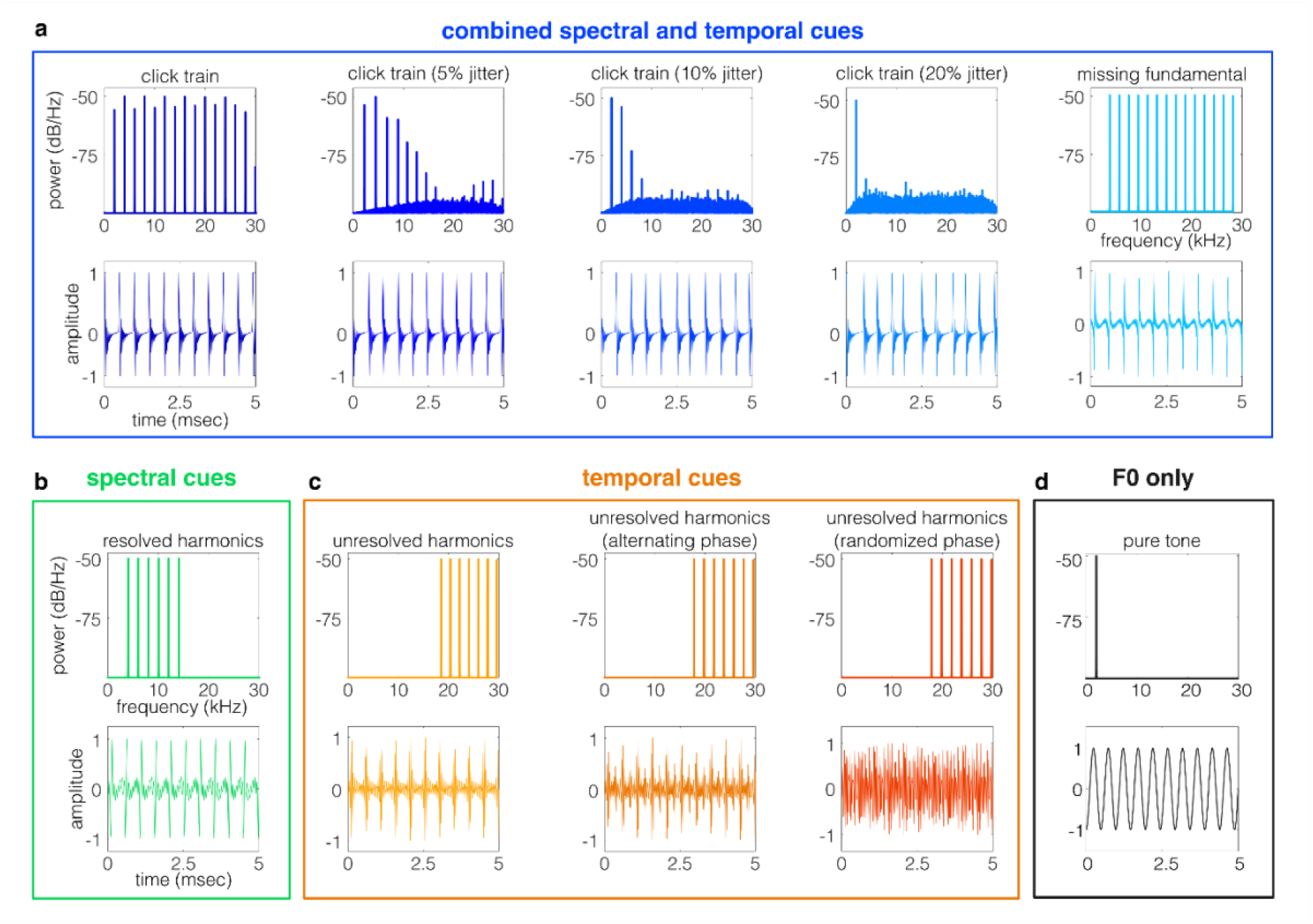
Stimuli. The spectrum (top plot) and temporal waveform (bottom plot) of all 10 timbres presented. Stimuli are coloured to indicate the 4 classes of F0 cue classes. **a**) Sounds with broad spectra contain both spectral and temporal pitch cues. Click trains contain energy at F0 and all harmonics, and their pitch cues can be parametrically weakened by randomly jittering the click timing to a greater degree (left to right). The missing fundamental sounds (right) contains energy at all harmonics except F0. **b**) Spectral pitch cues are contained in resolved harmonics (green). **c**) Unresolved harmonics (left) provide temporal pitch cues, which can be weakened through phase manipulations (middle and right). **d**) The pure tone contains energy only at F0.

The “*combined cues*” stimuli were broadband sounds that contained both resolved and unresolved harmonics up to 30 kHz, and therefore would provide both spectral and temporal F0 cues. The first timbre in this class was “click trains”, which were generated from temporally regular biphasic pulses at a rate of F0, producing all harmonics from F0 up to 30kHz (Figure 6a). We varied the temporal regularity of these click trains by temporally jittering the timing of the individual clicks by either 0% (perfectly periodic), 5%, 10%, or 20% of the inter-click interval. Stimuli with more jitter are less temporally regular, have less defined harmonic peaks in their spectra, and support poorer pitch discrimination in humans^32^ and ferrets (unpublished data from our lab). We also presented “missing fundamental” sounds, which contained all harmonics, both resolved and unresolved, but no energy at F0 (Figure 6a, right). The “*spectral cues”* stimulus contained only lower harmonics (Figure 6b) that are resolved in the ferret cochlea^33,70^. We presented three types of “*temporal cues”* stimuli (Figure 6c). All three contained the same set of higher, unresolved harmonics up to 30 kHz, but these frequency components were either: all in cosine phase (“unresolved harmonics”); in alternating sine and cosine phase, which doubles the periodicity of the sound (“alternating phase”); or in randomized phase, which diminishes the strong periodic peaks in the temporal waveform (“random phase”). Finally, we presented “pure tones”, which contained energy at F0 only (Figure 6d).

All sounds were presented via closed field, in-ear earphones (Panasonic RP-HV094E-K with an attached otoscope tip) at 70 dB SPL, with 25 ms onset and offset ramps. Stimuli were either 300 ms duration with 400 ms inter-stimulus silence interval (1 ferret), or 200 ms duration with 600 ms interstimulus intervals (3 ferrets). Each timbre and F0 combination was presented 13 times, in pseudo-random order.

At the beginning of each new electrode penetration, we presented 200 ms bursts of 30-60 dB SPL broadband Gaussian noise (300 ms interstimulus interval) to identify sound-responsive single units. Once responsive units were detected, a pseudorandomized sequence of pure tones (100 ms duration, 500 ms interstimulus interval, 40-80 dB SPL_A_ in 10 dB steps, 0.5-40 kHz in 0.45 octave steps) were presented to evaluate frequency tuning, allowing localization of every penetration within the tonotopic map. Finally, the complete set of pitch stimuli described above were presented.

### Spike sorting

Neural recordings were processed offline. Voltage traces were high-pass filtered at 150 Hz, and shared electrical noise across channels was removed with common average referencing. Spiking events were identified and clustered using the Kilosort2 algorithm (https://github.com/MouseLand/Kilosort2)^71^. Manual curation of the resulting spike clusters was carried out in Phy (https://github.com/cortex-lab/phy)^72^. Units exhibiting consistent spike waveforms with low variability and a distinct refractory period in their autocorrelograms were classified as single units. Only these well-isolated single units were retained for further analysis, and we refer to these as “neurons” throughout.

### Isolating sound-responsive neurons

We tested whether each neuron was responsive to the pooled set of pitch stimuli using a paired-sample t-test (α = 0.05). We compared the firing rate during the 150 ms of silence before the sound started with its firing rate during the 150 ms after sound onset, across all sound presentations. Response to sound offset was similarly tested in the 150 ms following sound offset. Only neurons that showed a significant response to sound onset and/or offset were included in further analysis.

### Decoding F0

Reponses from all sound-responsive neurons across the 18 electrode penetrations were used in the decoder, and trial order was randomly shuffled for each neuron independently to minimize effects of noise correlations in simultaneous recording. A trial was taken as the 400 ms after sound onset, and spikes were binned at 10 ms—the smallest bin size before decoding performance significantly worsened (Figure S3). For one animal, stimuli were 100 ms longer (300 ms instead of 200 ms), so neural responses 170-270 ms after onset were removed in this animal in order to align sound offset responses across animals. Each neuron’s firing rate was normalized by mean and variance across the whole data set by subtracting the neuron’s mean firing rate and dividing by the standard deviation in order to prevent decoding weights from reflecting each neuron’s baseline firing rate.

We used a linear model to decode F0 from a rank 3 tensor of neuronal activity, *X_nbt_*, where *n* is the neuron, *b* is the time bin in a trial, and *t* is the trial number (Figure 2a). The true F0 of each trial was defined as the target vector, *y_t_*. The decoder produced a prediction of each trial’s F0 from the sum of the element-wise product between the neuronal activity and a matrix of weights, *k*, with one weight per neuron and time point. The decoder’s predictions of each trial’s F0 is therefore defined as *ŷ_t_*, where:

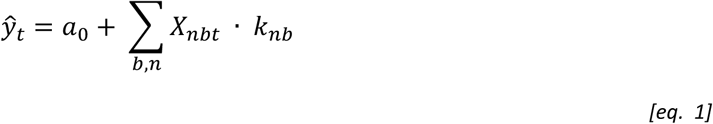

Model parameters were optimized with L2 regularization using glmnet (https://glmnet.stanford.edu/articles/glmnet.html) to minimize mean square error between *ŷ_t_* and *y_t_*. Trials were divided into 5 folds, each with 156 training trials and 39 non-overlapping test trials (with any leftover trials added to the test set of the fifth fold), and trials were randomly shuffled to promote an even sampling of F0s in each test set.

We began by training a separate decoding model for each of the 10 timbres presented. The fold with the lowest root mean square error (RMSE) between *ŷ_t_* and *y_t_* was taken as the fitted model for the given timbre. The decoding model for each timbre is therefore a 2-dimensional matrix with *N* rows and *B* columns, where *N* is the number of neurons and *B* is the number of time bins. Each value in this matrix is the coefficient for the activity of one neuron in one time bin, such that the sum of all values in the matrix multiplied by the neural activity on a single trial produces the estimation of that trial’s F0. Model weights can therefore be considered a proxy for F0 information.

We defined chance performance as the accuracy achieved when a null model was used, where all weights in the decoding kernel were zero. In this case, the decoder predicts the log median F0 of the series (1189 Hz) for every trial, producing an RMSE of 1.08.

A timbre-invariant version of the decoding model was run to examine whether F0 could be predicted across all stimulus timbres. The decoder predicted F0 using responses randomly drawn from all 10 timbres, while keeping the overall number of trials (and instances of each F0) in the training and test sets equal to the those in the single-timbre models above. We trained 1000 iterations of this model and report the best performing fold.

### Principal component analysis

We performed principal component analysis on the decoding kernels to provide a low-dimensional visualization of how F0 information evolved across response time and timbre types. The kernels for each of the 10 timbres were concatenated across time (without additional normalisation) to produce one matrix with *N* rows and (*B* ∗ *S*) columns, where *N* is the number of neurons, *B* is the number of time bins, and *S* is the number of stimuli (i.e. timbres). By applying PCA to this concatenated matrix, the dimensionality reduction isolated informative temporal spiking patterns that could be directly compared across the 10 timbres.

## Data Availability

Neural data have been deposited at Data Dryad as doi:10.5061/dryad.g79cnp64t, and are publicly available as of the date of publication. The code reported in this paper is available from the lead contact upon request.

## Supplementary Material

**Figure S1.**
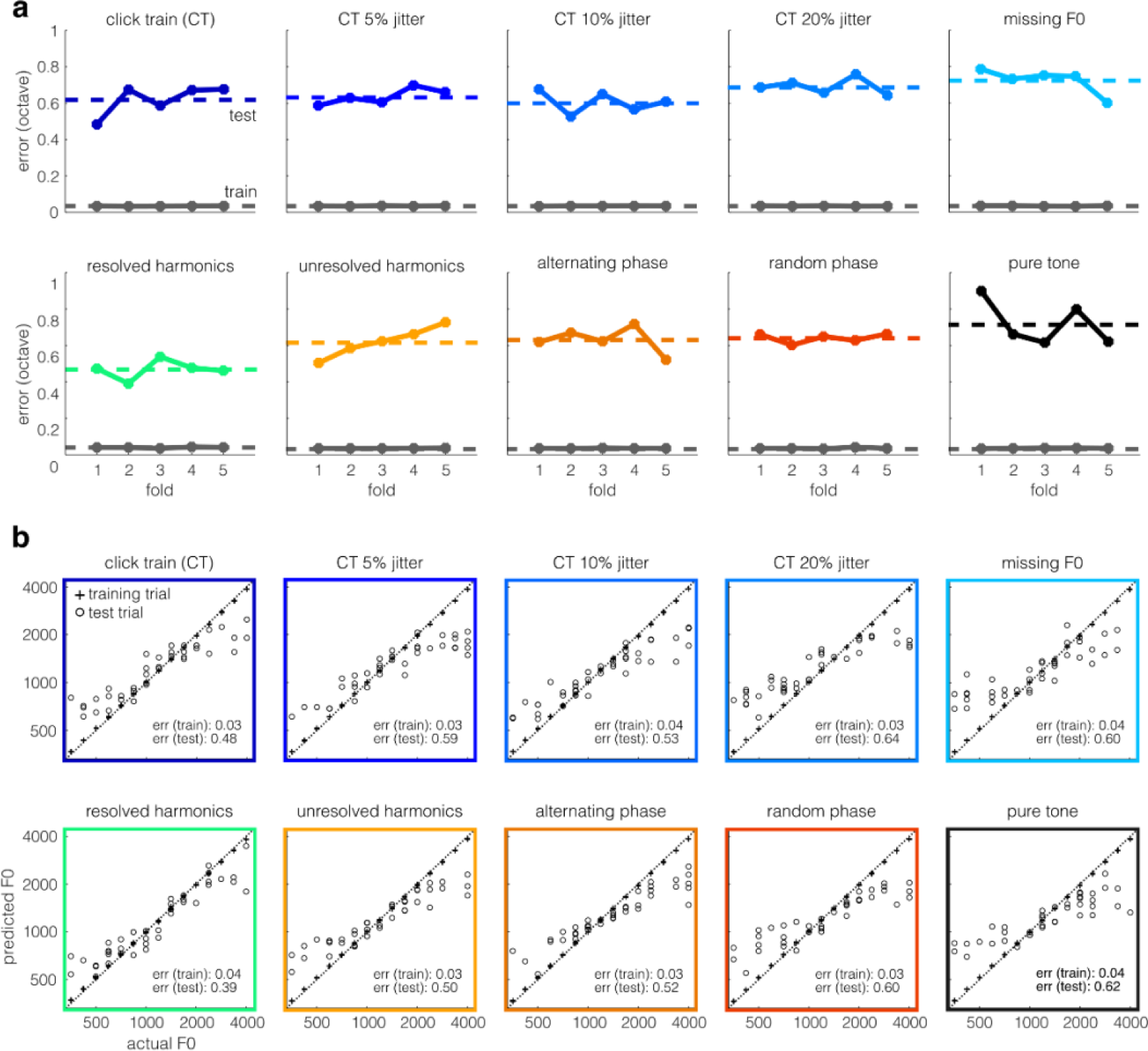
Decoding performance in all 5 training folds for each timbre. Related to Figure 2. **a**) Each plot shows the RMSE in octaves between the actual and predicted F0 across individual trials. Trials used in the training set are shown in grey, and the held-out test set is shown in colours corresponding to stimulus identity in Figure 6. The average RMSE across all 5 folds is shown as a dashed line. **b**) Each plot shows the decoding performance for individual trials in the training set (crosses) and test set (open circles) for the training fold with the smallest error (RMSE) for each timbre. Colours (blue, green, red, and black) indicate pitch cue classes described in Figure 6.

**Figure S2.**
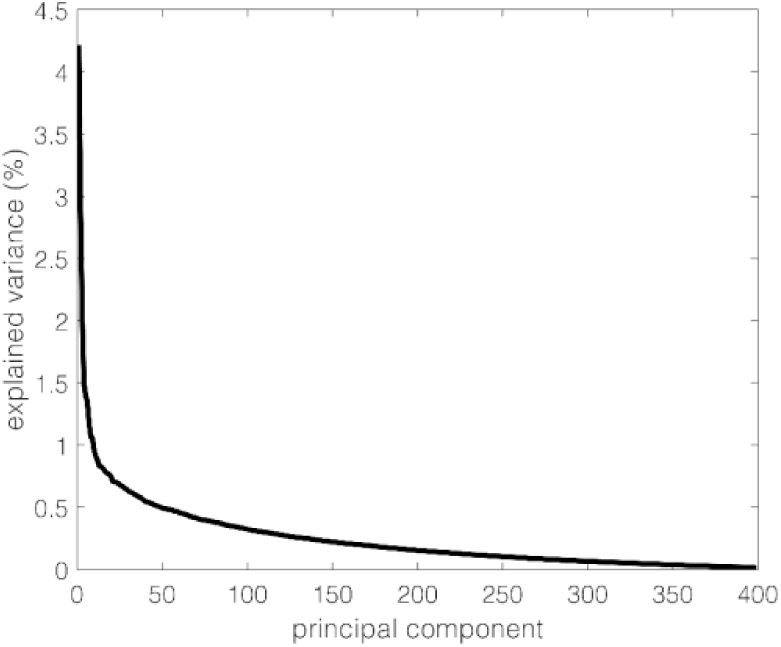
Scree plot of explained variance across principal components. Related to Figure 4.

**Figure S3.**
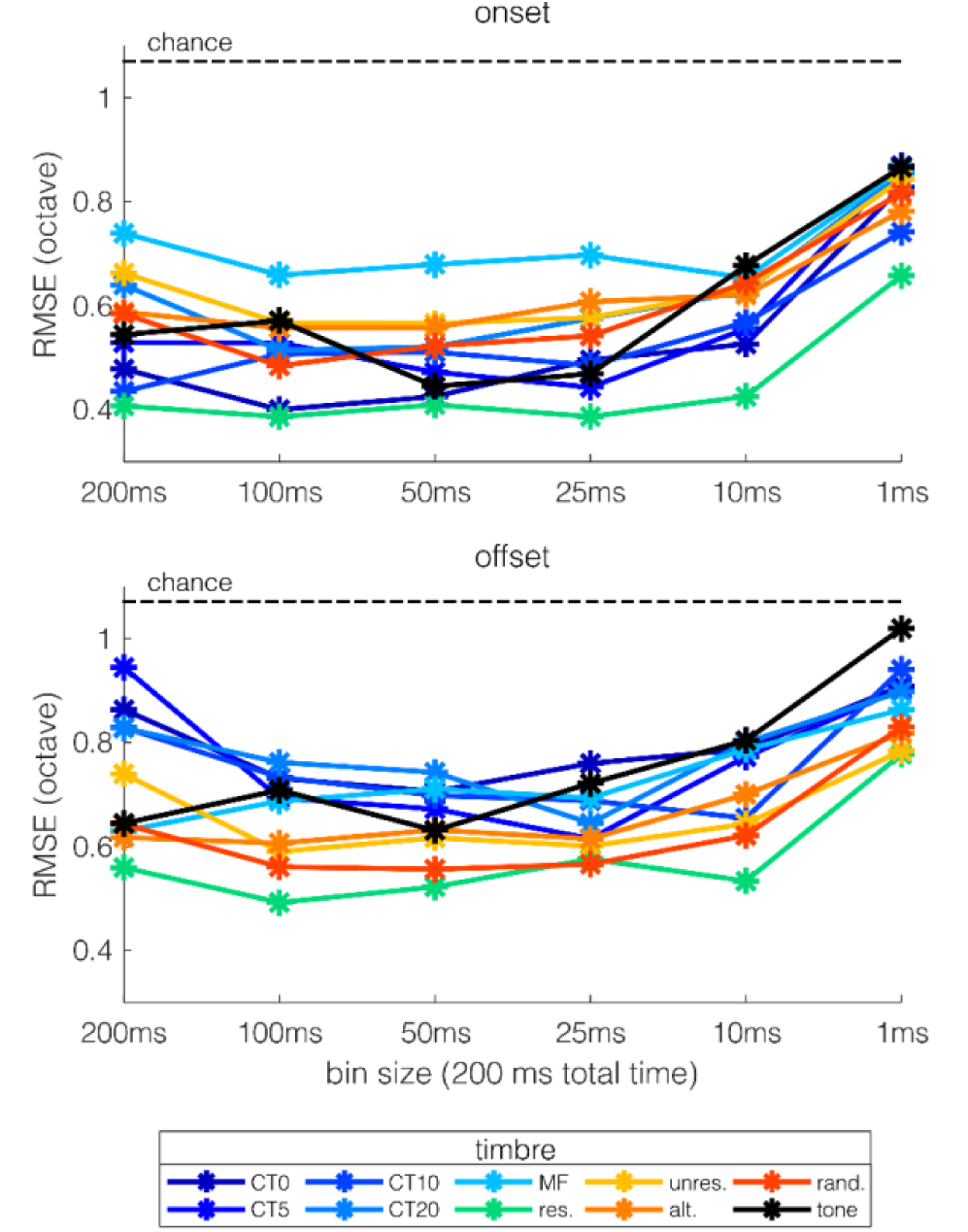
Decoding performance for different time bin sizes. Related to Figures 2 and 4. Decoders trained on neural activity in the first 200 ms after sound onset (top) perform similarly when spikes are binned at 200, 100, 50, 25, and 10 ms across all timbres (indicated by different colours, see Figure 6). Decoding performance is worse at 1 ms bins across all timbres. Decoders trained on neural responses to sound offset (bottom) perform similarly for bin sizes of 100, 50, 25, and 10 ms across all timbres, and are in general worse for bin sizes of 200 ms and 1 ms. Reduced accuracy for 1 ms bins is likely to due to greater inter-trial response variability at this temporal resolution, as well as excessive model parameters causing noisier fits for decoding weights.

**Figure S4.**
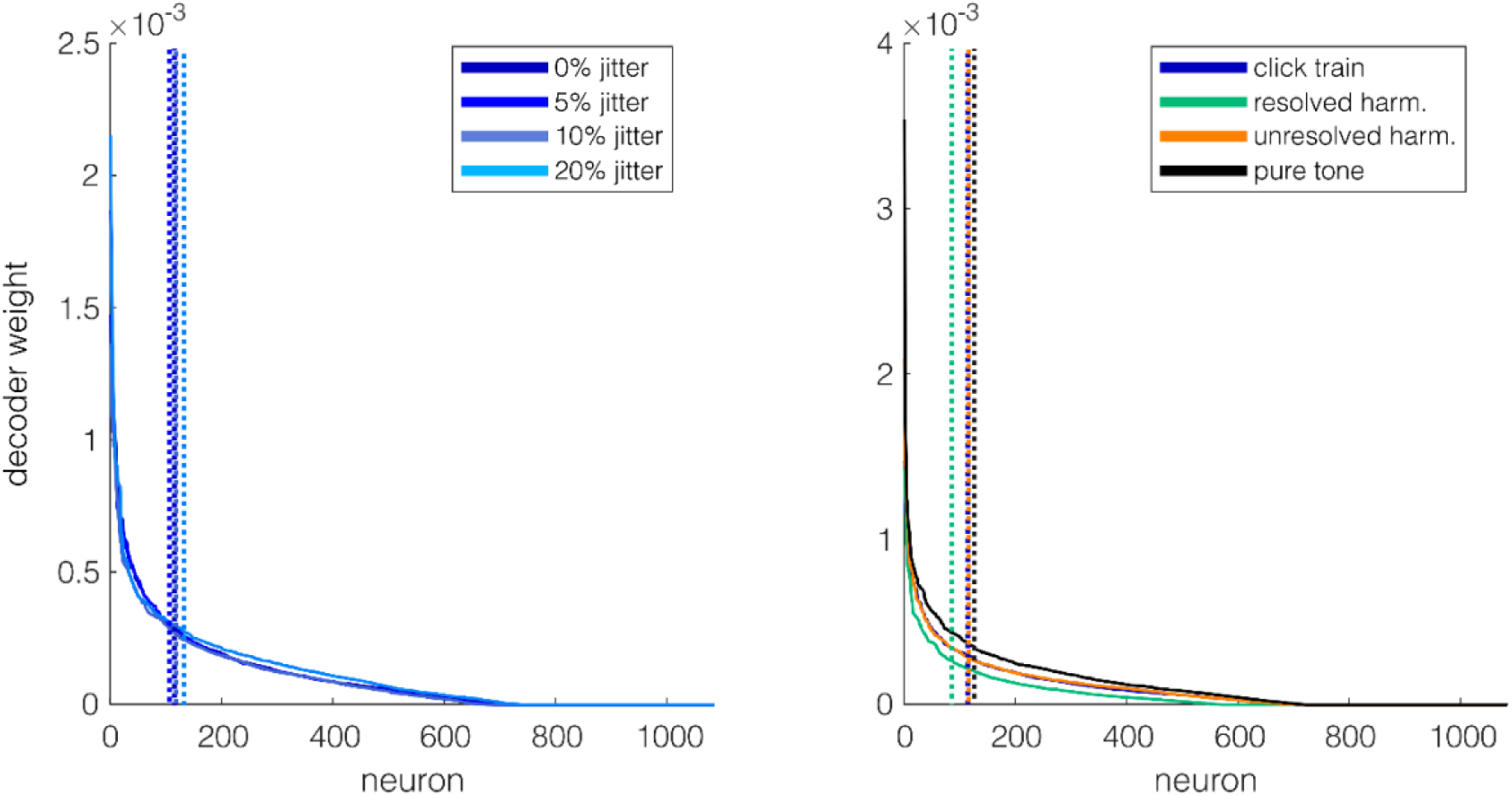
Distribution of the decoder weight magnitude across individual neurons. Related to Figure 5. For each timbre, neurons are ranked from largest to smallest summed absolute decoder weighting across time. Colours show timbre classes (left: click trains, right: representative stimuli from each pitch cue class). Dotted lines indicate 50% of the population’s total decoder weighting.

## Author contributions

VT analyzed the data and wrote the manuscript. QG collected the data. AJK edited the manuscript and secured funding. BW analyzed the data and edited the manuscript. KMMW conceptualized the experiment, collected the data, edited the manuscript, and secured funding.

## Acknowledgements

This work was funded by Biotechnology and Biological Sciences Research Council grants to K.M.M.W. (BB/M010929/1, BB/X013103/1), a Wellcome Trust Principal Research Fellowship to A.J.K. (WT108369/Z/2015/Z), and a University of Oxford Clarendon Scholarship to V.M.T.

## Materials & Correspondence

Kerry Walker

## References

1. Assmann, P. F. & Summerfield, Q. Perceptual segregation of concurrent vowels. The Journal of the Acoustical Society of America 82, S120 (1987).

2. Bird, J. & Darwin, C. J. Effects of a difference in fundamental frequency in separating two sentences. in Psychophysical and physiological advances in hearing (eds Palmer, A., Rees, A., Summerfield, A. Q. & Meddis, R.) 263–269 (Whurr, London, 1998).

3. de Cheveigné, A. Concurrent vowel identification. III. A neural model of harmonic interference cancellation. The Journal of the Acoustical Society of America 101, 2857–2865 (1997).

4. McPherson, M. J., Grace, R. C. & McDermott, J. H. Harmonicity aids hearing in noise. Attention, Perception, & Psychophysics 84, 1016–1042 (2022).

5. Arehart, K. H. Effects of harmonic content on complex-tone fundamental-frequency discrimination in hearing-impaired listeners. The Journal of the Acoustical Society of America 95, 3574–3585 (1994).

6. Hoekstra, A. & Ritsma, R. J. Perceptive hearing loss and frequency selectivity. in Psychophysics and Physiology of Hearing (eds Evans, E. F. & Wilson, J. P.) 263–271 (Academic Press, London, 1977).

7. McDermott, H. J. Music perception with cochlear implants: a review. Trends in Amplification 8, 49–82 (2004).

8. Walker, K. M. M., Bizley, J. K. & Schnupp, J. W. H. Cortical encoding of pitch: Recent results and open questions. Hearing Research 271, 74–87 (2011).

9. Moore, B. C. J. & Gockel, H. E. Resolvability of components in complex tones and implications for theories of pitch perception. Hearing Research 276, 88–97 (2011).

10. Cariani, P. A. & Delgutte, B. Neural correlates of the pitch of complex tones. I. Pitch and pitch salience. Journal of Neurophysiology 76, 1698–1716 (1996).

11. Carlyon, R. P. Encoding the fundamental frequency of a complex tone in the presence of a spectrally overlapping masker. The Journal of the Acoustical Society of America 99, 517–524 (1996).

12. Carlyon, R. P. Masker asynchrony impairs the fundamental-frequency discrimination of unresolved harmonics. The Journal of the Acoustical Society of America 99, 525–533 (1996).

13. Shackleton, T. M. & Carlyon, R. P. The role of resolved and unresolved harmonics in pitch perception and frequency modulation discrimination. The Journal of the Acoustical Society of America 95, 3529–3540 (1994).

14. Stewart, L., Von Kriegstein, K., Warren, J. D. & Griffiths, T. D. Music and the brain: disorders of musical listening. Brain 129, 2533–2553 (2006).

15. Tramo, M. J., Shah, G. D. & Braida, L. D. Functional Role of Auditory Cortex in Frequency Processing and Pitch Perception. Journal of Neurophysiology 87, 122–139 (2002).

16. Zatorre, R. J. Pitch perception of complex tones and human temporal-lobe function. The Journal of the Acoustical Society of America 84, 566–572 (1988).

17. Whitfield, I. C. Auditory cortex and the pitch of complex tones. The Journal of the Acoustical Society of America 67, 644–647 (1980).

18. Bendor, D., Osmanski, M. S. & Wang, X. Dual-Pitch Processing Mechanisms in Primate Auditory Cortex. Journal of Neuroscience 32, 16149–16161 (2012).

19. Bendor, D. & Wang, X. Neural Coding of Periodicity in Marmoset Auditory Cortex. Journal of Neurophysiology 103, 1809–1822 (2010).

20. Bendor, D. & Wang, X. The neuronal representation of pitch in primate auditory cortex. Nature 436, 1161–1165 (2005).

21. Tarka, V. M., Gaucher, Q. & Walker, K. M. M. Pitch selectivity in ferret auditory cortex. Current Biology 36, 3510–3522.e3 (2026).

22. Allen, E. J., Burton, P. C., Olman, C. A. & Oxenham, A. J. Representations of Pitch and Timbre Variation in Human Auditory Cortex. Journal of Neuroscience 37, 1284–1293 (2017).

23. Berger, J. I. et al. Distribution of multiunit pitch responses recorded intracranially from human auditory cortex. Cerebral Cortex 33, 9105–9116 (2023).

24. Bizley, J. K., Walker, K. M. M., King, A. J. & Schnupp, J. W. H. Neural Ensemble Codes for Stimulus Periodicity in Auditory Cortex. Journal of Neuroscience 30, 5078–5091 (2010).

25. Bizley, J. K., Walker, K. M. M., Silverman, B. W., King, A. J. & Schnupp, J. W. H. Interdependent Encoding of Pitch, Timbre, and Spatial Location in Auditory Cortex. Journal of Neuroscience 29, 2064–2075 (2009).

26. Fishman, Y. I., Micheyl, C. & Steinschneider, M. Neural Representation of Harmonic Complex Tones in Primary Auditory Cortex of the Awake Monkey. Journal of Neuroscience 33, 10312–10323 (2013).

27. Gander, P. E. et al. Direct electrophysiological mapping of human pitch-related processing in auditory cortex. Neuroimage 202, 116076 (2019).

28. Griffiths, T. D. et al. Direct recordings of pitch responses from human auditory cortex. Current Biology 20, 1128–1132 (2010).

29. Steinschneider, M., Reser, D. H., Fishman, Y. I., Schroeder, C. E. & Arezzo, J. C. Click train encoding in primary auditory cortex of the awake monkey: Evidence for two mechanisms subserving pitch perception. The Journal of the Acoustical Society of America 104, 2935–2955 (1998).

30. Walker, K. M. M., Bizley, J. K., King, A. J. & Schnupp, J. W. H. Multiplexed and robust representations of sound features in auditory cortex. The Journal of Neuroscience 31, 14565–14576 (2011).

31. Cunningham, J. P. & Yu, B. M. Dimensionality reduction for large-scale neural recordings. Nature Neuroscience 17, 1500–1509 (2014).

32. Pollack, I. Detection and Relative Discrimination of Auditory “Jitter”. The Journal of the Acoustical Society of America 43, 308–315 (1968).

33. Walker, K. M. M., Gonzalez, R., Kang, J. Z., McDermott, J. H. & King, A. J. Across-species differences in pitch perception are consistent with differences in cochlear filtering. eLife 8, e41626 (2019).

34. Fishman, Y. I. & Steinschneider, M. Temporally dynamic frequency tuning of population responses in monkey primary auditory cortex. Hearing Research 254, 64–76 (2009).

35. Carlyon, R. P. & Shackleton, T. M. Comparing the fundamental frequencies of resolved and unresolved harmonics: Evidence for two pitch mechanisms? The Journal of the Acoustical Society of America 95, 3541–3554 (1994).

36. Oxenham, A. J. Questions and controversies surrounding the perception and neural coding of pitch. Frontiers in Neuroscience 16, 1074752 (2023).

37. Klinge, A. & Klump, G. Mistuning detection and onset asynchrony in harmonic complexes in Mongolian gerbils. The Journal of the Acoustical Society of America 128, 280–290 (2010).

38. Osmanski, M. S., Song, X. & Wang, X. The Role of Harmonic Resolvability in Pitch Perception in a Vocal Nonhuman Primate, the Common Marmoset (*Callithrix jacchus*). The Journal of Neuroscience 33, 9161–9168 (2013).

39. Shofner, W. P. & Chaney, M. Processing pitch in a nonhuman mammal (Chinchilla laniger). Journal of Comparative Psychology 127, 142–153 (2013).

40. Hall, D. A. & Plack, C. J. Pitch Processing Sites in the Human Auditory Brain. Cerebral Cortex 19, 576–585 (2009).

41. Norman-Haignere, S., Kanwisher, N. & McDermott, J. H. Cortical pitch regions in humans respond primarily to resolved harmonics and are located in specific tonotopic regions of anterior auditory cortex. The Journal of Neuroscience 33, 19451–19469 (2013).

42. Bizley, J. K., Walker, K. M. M., Nodal, F. R., King, A. J. & Schnupp, J. W. H. Auditory cortex represents both pitch judgments and the corresponding acoustic cues. Current Biology 23, 620–625 (2013).

43. Qin, L., Chimoto, S., Sakai, M., Wang, J. & Sato, Y. Comparison between offset and onset responses of primary auditory cortex ON-OFF neurons in awake cats. Journal of Neurophysiology 97, 3421–3431 (2007).

44. Scholl, B., Gao, X. & Wehr, M. Nonoverlapping Sets of Synapses Drive On Responses and Off Responses in Auditory Cortex. Neuron 65, 412–421 (2010).

45. Sollini, J., Chapuis, G. A., Clopath, C. & Chadderton, P. ON-OFF receptive fields in auditory cortex diverge during development and contribute to directional sweep selectivity. Nature Communications 9, 2084 (2018).

46. He, J. On and off pathways segregated at the auditory thalamus of the guinea pig. The Journal of Neuroscience 21, 8672–8679 (2001).

47. Kopp-Scheinpflug, C., Sinclair, J. L. & Linden, J. F. When Sound Stops: Offset Responses in the Auditory System. Trends in Neurosciences 41, 712–728 (2018).

48. Campbell, R. A. A., Schulz, A. L., King, A. J. & Schnupp, J. W. H. Brief Sounds Evoke Prolonged Responses in Anesthetized Ferret Auditory Cortex. Journal of Neurophysiology 103, 2783–2793 (2010).

49. Hartley, D. E. H., Dahmen, J. C., King, A. J. & Schnupp, J. W. H. Binaural sensitivity changes between cortical on and off responses. Journal of Neurophysiology 106, 30–43 (2011).

50. Solyga, M. & Barkat, T. R. Emergence and function of cortical offset responses in sound termination detection. eLife 10, e72240 (2021).

51. Li, H. et al. Phasic Off responses of auditory cortex underlie perception of sound duration. Cell Reports 35, 109003 (2021).

52. Lee, J. & Rothschild, G. Encoding of acquired sound-sequence salience by auditory cortical offset responses. Cell Reports 37, 109927 (2021).

53. Lamothe, C., Bagur, S., Gosselin, E. & Bathellier, B. Sound offset responses become highly informative in the auditory cortex. The Journal of Physiology 604, 8702–8720 (2025).

54. Ali, F., Bury, G., Simon, A. & Linden, J. F. Distinctive neurophysiological correlates of sound onset and offset perception in humans. The Journal of Physiology 604, 8781–8795 (2026).

55. Bregman, A. S. Integration of Simultaneous Auditory Components. in Auditory Scene Analysis 213–394 (1990).

56. Colak, H. et al. Sound-offset encoding is related to speech-in-noise perception at sentence level in older adults. The Journal of Physiology 604, 8767–8780 (2026).

57. Dupret, D., Fusi, S. & Panzeri, S. Neural population activity for memory: Properties, computations, and codes. Neuron 114, 390–407 (2026).

58. Fishman, Y. I., Reser, D. H., Arezzo, J. C. & Steinschneider, M. Pitch vs. spectral encoding of harmonic complex tones in primary auditory cortex of the awake monkey. Brain Research 786, 18–30 (1998).

59. Kikuchi, Y. et al. The distribution and nature of responses to broadband sounds associated with pitch in the macaque auditory cortex. Cortex 120, 340–352 (2019).

60. Abrams, E. B., Marantz, A., Krementsov, I. & Gwilliams, L. Dynamics of Pitch Perception in the Auditory Cortex. Journal of Neuroscience 45, (2025).

61. Briley, P. M., Breakey, C. & Krumbholz, K. Evidence for Pitch Chroma Mapping in Human Auditory Cortex. Cerebral Cortex 23, 2601–2610 (2013).

62. Ince, R. A. A., Panzeri, S. & Kayser, C. Neural codes formed by small and temporally precise populations in auditory cortex. The Journal of Neuroscience 33, 18277–18287 (2013).

63. Kampa, B. M., Roth, M. M., Göbel, W. & Helmchen, F. Representation of visual scenes by local neuronal populations in layer 2/3 of mouse visual cortex. Frontiers in Neural Circuits 5, 18 (2011).

64. Resulaj, A., Ruediger, S., Olsen, S. R. & Scanziani, M. First spikes in visual cortex enable perceptual discrimination. Elife 7, e34044 (2018).

65. Ben Hamed, S., Schieber, M. H. & Pouget, A. Decoding M1 neurons during multiple finger movements. Journal of Neurophysiology 98, 327–333 (2007).

66. Rickert, J., Riehle, A., Aertsen, A., Rotter, S. & Nawrot, M. P. Dynamic encoding of movement direction in motor cortical neurons. The Journal of Neuroscience 29, 13870–13882 (2009).

67. Averbeck, B. B., Latham, P. E. & Pouget, A. Neural correlations, population coding and computation. Nature Reviews Neuroscience 7, 358–366 (2006).

68. Jun, J. J. et al. Fully integrated silicon probes for high-density recording of neural activity. Nature 551, 232–236 (2017).

69. Karsh, B. SpikeGLX. Janelia Research Campus (2017).

70. Sumner, C. J. & Palmer, A. R. Auditory nerve fibre responses in the ferret. European Journal of Neuroscience 36, 2428–2439 (2012).

71. Pachitariu, M., Sridhar, S., Pennington, J. & Stringer, C. Spike sorting with Kilosort4. Nature Methods 21, 914–921 (2024).

72. Rossant, C. et al. Spike sorting for large, dense electrode arrays. Nature Neuroscience 19, 634–641 (2016).

